# Fitness and mating consequences of variation in male allocation in a wind pollinated plant

**DOI:** 10.1101/2021.12.01.470797

**Authors:** Abrar A. Aljiboury, Jannice Friedman

## Abstract

In hermaphrodites, the allocation of resources to each sex function can influence fitness through reproductive success and mating success. In wind pollinated plants, sex allocation theory predicts that male fitness increases linearly with investment of resources into male function. However, there have been few empirical tests of this prediction. We experimentally manipulated allocation to male function in *Ambrosia artemisiifolia* (common ragweed) in a field experiment and measured mating success using genetic assays. We investigated the effects of various morphological traits and flowering phenology on male fitness, and on male and female mate diversity. Our results provide evidence for a linear relation between allocation to male function and fitness. We find earlier onset of male flowering time increases reproductive success, while later onset flowering time increases the probability of mating with diverse individuals. This research is among the first empirical studies testing the prediction of linear male fitness returns in wind pollinated plants. Our results provide insight into the large investment into male function by wind pollinated plants and temporal variation in sex allocation.

## Introduction

Hermaphroditic organisms attain fitness through both female and male sex functions and they can maximize reproductive success through a wide variety of different strategies. Most flowering plants are hermaphroditic and because of vector-mediated gamete transfer, plant mating can be highly promiscuous with individuals mating with numerous sexual partners through either sex function. The ecological and genetic factors that determine reproductive success have been well documented for female function because of the relative ease of assessing fitness through maternal contribution. In contrast, success through male function has been more difficult to quantify because it relies on using molecular markers to identify paternity. With current molecular techniques, we are beginning to gain an understanding of male mating success, including who has mated with whom and how often (e.g. Tomaszewski et al. 2018; Christopher et al. 2019; Santos del Blanco et al. 2019).

Sex allocation theory considers how the fitness acquired through each sex function varies with the investment of limited resources into female or male reproductive structures (Charnov 1982; Charlesworth 1991; Brunet 1992; Emms 1993). In general, hermaphroditic organisms should invest in both sex functions to the point where the marginal fitness returns are equal, at which point investment should turn to the sex function with the more linear, or less decelerating, sex function (Charlesworth 1991; Brunet 1992). The shape of the fitness gain curve is determined by a variety of intrinsic and extrinsic factors, including dispersal dynamics, biotic interactions, sibling competition, and mating patterns (Harder and Thomson 1989; Harder and Barrett 1995; Zhang 2006). While there has been extensive theoretical treatments of sex allocation theory, empirical support for their predictions, particularly regarding fitness through male function, remains limited.

In sessile plants that rely on external vectors for transferring and receiving pollen, various features of the pollination environment can affect the shapes of fitness gain curves. Many animal pollinated plants likely have diminishing returns on investment in male function due to a variety of processes, including pollinators becoming saturated with pollen after visiting a plant (Lloyd 1984), pollinator grooming removing pollen (Harder 1990), and deposition of related pollen causing local mate competition (Charnov 1982). In contrast, in wind pollinated plants the male gain curve is expected to be more linear (Burd and Allen 1988; Klinkhamer et al. 1997; Sakai and Sakai 2003). There are several explanations for this. First, it is unlikely that the air can become saturated with pollen, at least within the scope of biologically plausible pollen production. Second, once pollen is liberated from a plant, pollen loss from the airstream should be stochastic and independent of an individual plant (Niklas 1985). And finally, conspecific pollen captured by stigmas is likely proportional to its concentration in the air (although there might be subtle differences depending on pollen size (Paw U and Hotton 1989; Friedman and Harder 2005). Similar arguments have been made for the male gain curve in sessile spermcasting marine organisms (reviewed in Schärer 2009). Together these factors suggest that for a given individual, producing more pollen should result in siring more offspring (ie. linear relation between allocation to pollen production and fitness).

Sex allocation theory also explains shifts in relative resource allocation to female or male function with plant size (Klinkhamer et al. 1997; Zhang 2006). Larger and taller plants often have more resources to invest in gamete production leading to greater fecundity. This can produce a “budget” effect of plant size on sex allocation. Generally, access to more resources is associated with greater relative allocation to female function (Lloyd 1984; Korpelainen 1998; Chen et al. 2017), because of the greater cost associated with producing seeds and fruit. In addition to the budget effect, size can also have a direct effect on fitness, where larger plants have greater fitness than small plants for a given investment in reproduction. For example, in wind pollinated plants, the physical placement of male flowers at higher positions or at the tips of long branches, results in more effective pollen dispersal and greater siring success (Young and Schmitt 1995; Tonnabel et al. 2019a). The influence of budget and direct effects of plant size on sex function may be aligned, but the underlying mechanism might be through selection on fecundity or through access to more mates.

Male mating with more than one female partner is almost ubiquitous in the flowering plants although whether this arises as a by-product of selection on siring success in general, or on outcross mate diversity in particular is not clear (Pannell and Labouche 2013). Mating with multiple partners can provide an advantage— regardless of any increase in total fecundity—by increasing the genetic diversity of offspring (Barrett and Harder 2017). Higher mate diversity almost certainly increases the variance in offspring genotypes and may decrease the variance in final offspring number (for example, by minimizing the risk of unsuccessful pairings) which raises the probability of successfully leaving offspring (Gillespie 1974; 1977). Producing a set of genetically diverse offspring increases the probability of generating a “winning” phenotype (Williams 1975; Maynard Smith 1976). Increasing the variance in offspring genotypes is especially beneficial in heterogeneous environments if it produces genotypes that succeed in different conditions, akin to bet-hedging strategies (Antonovics and Ellstrand 1984; Simons 2011). Finally, greater mate diversity may reduce sibling competition between related sibs (Karron and Marshall 1990). From the maternal perspective, a common measure of mate diversity is to determine the genetic contribution of different fathers to the seeds on a plant (correlated paternity, *r*_p_: Ritland 2002). This measure can be extended to consider the entire mating portfolio of a plant (Barrett and Harder 2017) through both male and female function, although a full quantification of mate diversity is rare (but see Tomaszewski et al. 2018; Christopher et al. 2019).

Mating opportunities between plants are influenced by the timing of female and male function (Lloyd 1980). The vast majority of studies on the evolution of flowering time have focused on animal pollinated plants, and fitness through female function (Christopher et al. 2020), but of course selection through the two sex functions need not be in harmony (Delph and Ashman 2006). Selection on male flowering time may be driven by mating opportunity or through genetic covariation with other traits (Austen and Weis 2016a). Furthermore, many hermaphroditic and monecious plants have some temporal separation in the onset of female and male function (dichogamy; Bertin and Newman 1993) that leads to a shift in the mating environment (the relative abundance of female and male phase flowers) through time. For example, in protandrous plants, the floral sex ratio shifts from male- to female-dominated during a population’s flowering season (Brunet and Charlesworth 1995; Brookes and Jesson 2010). This means that in general, early male-phase flowers encounter pollen competition for few ovules compared with later male-phase flowers that have access to more ovules (Nakamura et al. 1989; Stanton 1994). This effect is weakened if protandry in incomplete and flowering duration is long with substantial overlap in sex function.

Our primary goal in this study is to use a manipulative field experiment to evaluate the effect of allocation to male function on siring success and mate diversity in a wind pollinated herbaceous plant. We also set out to determine if there were additional benefits accrued through plant size (height, width, and plant biomass) on siring success; and to determine the effect of flowering time on mating opportunities and reproductive success through male function. We used the monoecious herb, *Ambrosia artemisiifolia* (common ragweed), a weedy annual plant that is known to produce a prodigious amount of allergenic pollen. It has been well established in *A. artemisiifolia* that sex allocation is both size and resource dependent, where taller plants and those with high light availability invest more in male function (McKone and Tonkyn 1986; Ackerly and Jasieński 1990; Traveset, 1992; Paquin and Aarssen 2004; Friedman and Barrett 2011a; Nakahara et al. 2018). To achieve our aims we experimentally manipulated allocation to male function to limit confounding sex allocation with overall condition or plant size and to produce individuals with a range of different male allocation patterns (Emms 1993, Schärer 2009). After allowing plants to naturally wind pollinate, we used genetic markers to estimate the progeny sired by each plant. This study represents the first attempt to estimate the male gain curve in a wind-pollinated plant after experimentally controlling for male allocation (similar to a series of studies in free-spawning animals; Yund and McCartney 1994, McCartney 1997, Yund 1998, Johnson and Yund 2009) and to quantify mate diversity through paternal function.

## Methods

### Study Species

*Ambrosia artemisiifolia* L. (common ragweed) is an early successional weed that inhabits a broad range of habitats including roadsides, cultivated fields, and disturbed lands (Bazzaz 1974). The species is monoecious, producing male inflorescences at the tips of branches and inconspicuous female flowers in leaf axils (Payne 1963). Ragweed can produce between 200 and 6000 seeds depending on the plant’s condition (Fumanal et al. 2007), although fecundity is substantially lower when plants are disturbed or transplanted. Plants are usually weakly protandrous at the plant level (i.e. male flowers open before female flowers) and there is plasticity in the degree and order of dichogamy (Friedman and Barrett 2011a). Ragweed is self-incompatible (Friedman and Barrett 2008), so that all seed are outcrossed. In addition, it is an annual plant reproducing entirely through seed (Bassett and Crompton 1975), which allows for more easily quantifying fitness.

### Experimental Design

Between June 1-5 2017, we collected *A. artemisiifolia* seedlings at the 4-leaf stage from 18 natural populations around Syracuse, NY, USA. The populations were selected arbitrarily, ensuring they were approximately 4 km apart (range: 4-9 km). We collected plants from multiple sites to increase the genetic variation in microsatellite loci, which we would later use for paternity assignment. We dug up seedlings and immediately transplanted them into 2.5 cm pots filled with Sunshine Mix4 (Sun Gro Horticulture). We maintained the plants in their pots for 8 days, then transplanted them into randomly predetermined locations in three blocks in a field at Syracuse University, NY, USA (43°00’47’N 76°07’07’W). The three blocks were spaced approximately 15m from each other, and each block was covered with groundcloth to inhibit weed growth. Within each block, we planted 64 plants in a square grid, with individuals 50cm away from each other. We planted equal numbers of all subpopulations into each block (n=4 per subpopulation per block) and recorded the specific location of each plant. We monitored the plants’ condition and replaced any dead plants up to two weeks after transplanting in the field (n=12 transplants).

We minimized pollen contamination by naturally occurring ragweed by mowing the field around the three blocks every two weeks and manually removing any ragweed surrounding the field site at least twice a week. The larger area around the experimental garden was maintained by Syracuse University and received considerable mowing and horticultural maintenance, which reduced the overall presence of ragweed plants.

To limit confounding sex allocation with overall condition or plant size (Emms 1993, Schärer 2009), we artificially manipulated male allocation by randomly assigning each plant to one of four male allocation categories that differed in their maximum number of branches bearing male inflorescences per plant. We started the manipulations between August 1-5 (which coincided with when the very first plants began flowering) after allowing plants to grow for about seven weeks. We cut male inflorescences from each plant to match its pre-assigned allocation category, and maintained the number of inflorescences by removing a subset of newly emerged male inflorescences twice a week. The four categories had a maximum of 4, 8, 16, or 32 branches bearing male inflorescences per plant. Although the number of branches increased exponentially, subsequent branches have fewer and shorter inflorescences, so this approximated a linear increase in male flowers (Figure S1: category 4 has greater variance because we could only remove excess branches from the randomly assigned individuals).

We measured plant height from ground level to the base of the inflorescence spike, and measured width at the widest point of the plant including branches. We recorded these size measurements three weeks after transplanting (17 July 2017) and repeated the measurements on 7 August and 7 September. We also recorded the date the first female and male flower opened on each plant.

Subsequent to the initial difference in onset of female and male flowers, plants continue producing new flowers of both sex functions throughout the blooming period (see Figure 4 in Friedman and Barrett 2011a). Once plants started senescing (between October 2 and 15, 2017), we harvested male inflorescences from each plant and separately collected above-ground plant material. We dried the male inflorescences and above-ground plant material for 3 days at 70°C, then recorded the biomass as proxies of male allocation and plant size, respectively.

We allowed plants to pollinate naturally within the field. We randomly selected a subset of plants (n=24 per plot, total=72) from which we collected at least 60 seeds twice: once when the plants first matured seeds (between September 12 and 26, 2017), and once before the plants were harvested (between October 1 and 12, 2017). We refer to these plants as ‘maternal’ plants but recall that the species is monoecious, so these focal 24 plants are also ‘paternal’ plants. We ensured that the maternal plants were evenly represented among the four male allocation categories. We avoided collecting seeds from plants at the edges of plots as they were not surrounded by experimental plants in all directions.

### Parent and progeny genotyping

We collected fresh leaf tissue from all experimental plants in the field on August 23-25 for subsequent genotyping. We stored leaf tissue at -80°C until DNA extractions could be performed. On December 21, 2017, we stratified 60 seeds (from an equal mix of early and late collection dates) from each of the 72 focal maternal plants in the dark at 4°C for 16 weeks, and then planted them in 96-well plug trays filled with Sunshine Mix4 mixed with 25% sand. We planted more seeds than we intended to genotype to ensure we had 24 seedlings per maternal plant. The average germination probability of the seeds was 68.5 percent. We maintained plants in the greenhouse under 16-hr photoperiod and 21°C/18°C day/night temperatures. After six weeks, once the majority of plants were at the 4-leaf stage or larger, we collected fresh leaf tissue from 24 progeny per maternal plant, and stored the plates at -80°C until DNA extractions could be performed.

We extracted DNA from frozen leaf tissue of the 1728 progeny and 192 field plants using a modified CTAB protocol. We then used seven polymorphic microsatellite markers that were previously developed for *A. artemisiifolia* (Genton et al. 2005; Chun et al. 2009) to identify male parentage. We performed polymerase chain reaction using Bio-Rad thermocyclers (see Table S1) and labelled PCR products were analyzed at the Institute of Biotechnology at Cornell University on an ABI 3730xl DNA Analyzer using GeneScan LIZ500 size standard (Applied Biosystems, Foster City USA). We manually scored the genotypes using GeneMarker v.2.2.0 (Soft Genetics, State College USA). We used CERVUS version 3.0.7 (Field Genetics Ltd, London UK), to perform paternity analyses using stringent parameters with a minimum of 3 typed loci and accounting for a 1% genotyping error rate and assigned paternity using a strict (95%) confidence criterion.

### Data analysis

We assessed reproductive success in three ways. First, we quantified the total number of seeds sired by each plant and refer to this as *male reproductive success*. Second, we quantified the number of unique plants on which a given individual sired seed and we refer to this as *male mate diversity*. Third, we quantified the number of seeds sired by different pollen donor plants on a given focal plant and we refer to this as *female mate diversity*. Note that we did not quantify total seed production by plants (female reproductive success). Our primary aim was to evaluate the association between male allocation and the measures of male reproductive success (total number of seeds sired and number of mates), including their relation with each other, and effects of size and flowering time.

All analyses were performed using R version 4.1.1 (R Core Team 2021). We used generalized linear mixed models (GLMMs) using the glmmTMB package (Brooks et al. 2017) and marginal means were estimated using the package emmeans package (Lenth 2021) We used the DHARMa package for diagnostic tests of residuals (Hartig 2017). We had three sets of models, where the response variable was either male reproductive success, male mate diversity, or female mate diversity. In all models, we included the fixed effects of male allocation, female flowering day, male flowering day, plant width, plant height, biomass, and whether the plant was located on the edge or interior. The latter effect was excluded in models of female mate diversity since all plants were located in the interior; in this model we additionally included germination probability of seed. Two random effects were included in all models: block and source population. We identified significant terms in the model using AIC scores and log-likelihood ratios of model fit, and present the simplest model of best fit (except when the term reflected experimental design, like edge effect). We tested for significant differences among categories using Dunn-Sidak adjustment for multiple testing. Recall that we quantified male allocation in two ways—the categorical manipulation the plant received, and the continuous measure of male inflorescence weight. Thus, we ran separate models including either measurement as a fixed effect. The two sets of models produced very similar results, which is not surprising because the two measures of male allocation are strongly statistically associated (Figure S1).

Models of male reproductive success used a negative binomial distribution with the canonical link function. Additionally, to specifically evaluate the shape of the male gain curve, we fit a power function (*y*=*ax^b^*; Charnov 1979; Johnson and Yund 2009) to the continuous male allocation data. The exponent of the power function (*b*) describes the shape of the curve. If the exponent does not differ significantly from 1 then reproductive success approximates a linear function of male allocation; if it is significantly <1 or >1, reproductive success is either a saturating or accelerating function of male allocation, respectively. Models of mate diversity required accounting for the number of seeds sired. Thus, we used a binomial model (unique mates/total seeds sired) with prior weights, which best captured the underlying biological process and accommodated the frequent occurrence of the upper bound. Models of female diversity used a poisson distribution with an offset parameter of the number of seeds genotyped and the canonical link function.

We investigated the fitness accrued through male function by mating with additional mates by estimating the slope of a least-square regression between standardized male mating success and standardized male reproductive success (‘Bateman gradient’: Arnold and Wade 1984). The model included covariates for flowering time, male allocation, edge location, and random effects accounting for block and source population. We compared the likelihood of linear and quadratic terms in the regression using AIC scores and log-likelihood ratios of model fit.

We considered the spatial dispersal of successful pollen transfer by calculating the Euclidean distance between a pollen donor and the plant on which it sired offspring. Note that because of our experimental design there was a 15m gap between any two blocks. We modeled mating distance using generalized linear mixed models (GLMMs) using the glmmTMB package, with a gamma distribution. To investigate the effect of morphological and phenological variables on dispersal distance, we included male inflorescence weight, plant height, plant width, biomass, female flowering time, male flowering time and edge as fixed effects, and candidate father identity as a repeated random effect. To investigate the relation between mean dispersal distance and reproductive success we modeled the effects of male siring success and male mate diversity as fixed effects. To visualize variation in dispersal kernels, for every pollen donor plant that sired more than 5 seeds, we fit a Weibull distribution to its dispersal kernel, and extracted the shape parameter, *κ*, and the scale parameter, *λ* for each pollen donor plant. We also used the software NM*π* (Chybicki 2018) to calculate the parameters of the pollen dispersal model. Specifically, we estimated average dispersal distance (*δ*_p_), the shape of the exponential-power dispersal kernel (*b*_p_), the intensity of directionality of dispersal (*κ*_p_), and the prevailing direction of pollen dispersal (*α*_p_).

## Results

### Identification of mating outcomes

A total of 1257 (72.7%) offspring were assigned paternity, the remaining 471 (27.3%) individuals were excluded because either confidence was low, or no paternity matches were found using the parental genotypes (ie. the seed were sired by plants outside the experimental fields). Of the 192 experimental plants, 176 plants (91.7%) sired seed. Of the 16 plants that were not represented in the paternity of seed, three plants were removed from the analysis because they were typed at less than three loci, and the remaining 13 plants were not identified as the father for any genotyped seed.

### Pollen dispersal and siring distance

The incidence of mating strongly declined with distance, with a heavy-tailed distribution. Plants sired most of their seeds locally, where siring peaked within half a meter of pollen source and declined rapidly over the 25 m distance of the field (distance between mates: mean = 2.81 m, SD = 5.03 m.; Figure 1A, solid line). Known intermate distances were vastly smaller than the distances between all pairs of plants (ie. all possible mates; *t*=52.91, *P*<0.001) and also smaller than the distances of all within-block pairs of plants (*t*=4.39, *P*<0.001) —reflecting the predominance of mating amongst near neighbours. Of the seed we successfully genotyped, 90.7% were sired by fathers within the same experimental block (1127 of 1242), and only 9.3% were due to longer distance siring events between blocks. None of the morphological or phenological variables we investigated (male inflorescence weight (or allocation category), plant biomass, width, height, the onset of female or male flowering) had any effect on siring distance, and there was also no relation between siring distance and either the number of seeds sired or mate diversity. The only variable that was significantly associated with dispersal distance was whether plants were located in the interior or the edge (adjusted for other effects in the model, mean dispersal distance: edge: 2.92 m, 95% CI: 2.38-3.59; interior: 2.24 m, 95% CI: 1.89-2.66; *t*=1.94, *P*<0.05).

**Figure 1.**
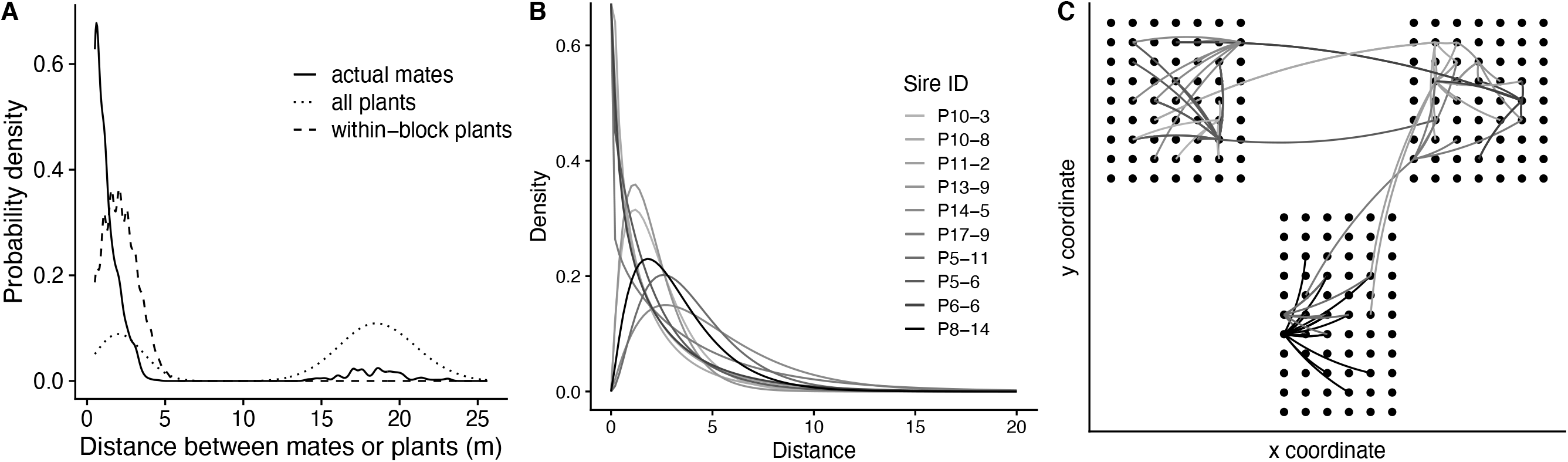
(A) The frequency distributions of interplant and intermate distances of *Ambrosia artemiisfolia* plants in experimental arrays. The solid line represents the probability density distribution of distance between mates in our progeny assays, while the dotted and dashed lines represent the probability density distributions of the distance between all plants in the experiment (dotted line) and distances between plants within a block (dashed line). (B) Modelled Weibull distribution of the dispersal kernel for a random subset of 10 experimental plants. (C) Distribution of 192 plants in experimental arrays (black dots) with the curves showing the trajectory of pollen to reach the known mates for the same random subset of 10 experimental plants. Note that the physical distance between the three blocks has been compressed for ease of figure presentation.

There was substantial variation among plants in their fitted dispersal distributions. To visualize the variation in the distribution of inter-mate distances, we plot the fitted Weibull distribution for a random subset of 10 plants (Figure 1B) and also show the trajectories of pollen dispersal for the same subset of 10 plants (Figure 1C). The Weibull distribution is described by the shape parameter, *κ*, and the scale parameter, *λ*. Among individual candidate fathers, estimated shape *κ* had range=0.62-4.74 (mean=1.51) and the estimated scale *λ* had range=0.72-6.39 (mean=2.30). We next used NM*π* (Chybicki 2018) to calculate properties of the pollen dispersal kernel.

Similar to the results using other methods, the estimated average dispersal distance (*δ*_p_) was 1.2m with high variance (S.E.=1.3m), the shape of the exponential-power dispersal kernel (*b*_p_) was 0.18 (S.E.=0.14), the intensity of directionality of dispersal (*κ*_p_) was 0.48 (S.E.=0.31), and the prevailing direction of pollen dispersal (*α*_p_) was 2.68 with high variance (S.E.=8.93).

### Male reproductive success

Individual plants sired a mean=6.60 (SD=6.53, Range=0-39) seed, of the successfully genotyped offspring. The effect of our experimental manipulations resulted in plants with greater male allocation siring significantly more seed than plants with less male allocation (Table 1A; Figure 2A). Adjusted for other effects in the model, plants in the lowest allocation category sired an average of 4.44 seeds (95% CI: 3.36-5.87) while plants in the highest allocation category sired an average of 8.05 seeds (95% CI: 6.28-10.31). When we consider allocation as a continuous variable modeled as male dry inflorescence weight, we found that the number of offspring sired increases significantly with male inflorescence weight (Table 1B; Figure 2B). The exponent of the power function (*b*=0.85, SE=0.30) saturates slightly, but did not significantly differ from one (*t*_168_= -0.45, *P*=0.65), indicating a significant linear relation between male allocation and siring success (*F*_1,168_=7.68; *P*<0.001).

**Table 1.** Models of effects on male reproductive success in Ambrosia artemiisifolia. Results for A) model with factorial measure of male allocation (manipulation category), and B) model with continuous measure of male allocation (male inflorescence weight), using negative binomial GLMMs. Random effects (not shown) included the effects of source population and block. Male flowering time is abbreviated MFT, and Male inflorescence weight is abbreviated MIW. Bold P-values indicate significant effects.

|  | **Term Condition** | **df** | **$\chi^2$** | ***P*-value** |
| --- | --- | --- | --- | --- |
| A) | Allocation category | 3 | 9.03 | **<0.05** |
|  | Male flowering time | 1 | 9.34 | **<0.001** |
|  | Plant width | 1 | 4.44 | **<0.05** |
|  | Edge | 1 | 5.52 | **<0.05** |
|  | Allocation x MFT | 3 | 9.46 | **<0.05** |
| B) | Male inflorescence weight | 1 | 6.67 | **<0.01** |
|  | Male flowering time | 1 | 13.04 | **<0.001** |
|  | Plant width | 1 | 5.99 | **<0.05** |
|  | Edge | 1 | 5.62 | **<0.05** |
|  | MIW x MFT | 1 | 7.23 | **<0.01** |

**Figure 2.**
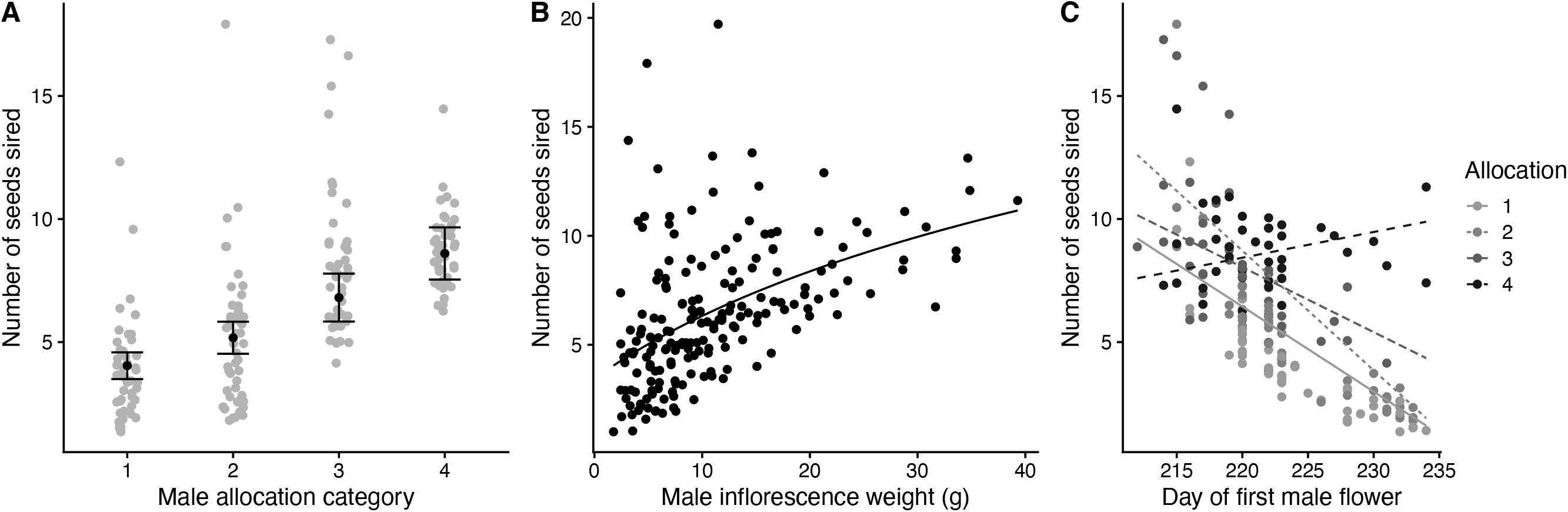
Influences on the number of seeds sired (male reproductive success) in *Ambrosia artemiisfolia* plants, including (A) treatment effects of four experimentally manipulated categories of male allocation, (B) male inflorescence weight, and (C) interaction between male allocation category and the julian day of the start of male flowering. Panel B shows the relation described by the power function regression, *y*=*ax^b^,* with *b*=0.85. All results represent model- adjusted and back-transformed values from ln estimates. See Table 1 for statistical details.

We found a strong effect of the timing of onset of male flowering on male reproductive success where earlier flowering plants sired significantly more seed (Table 1). Moreover, we found a significant interaction between allocation and male flowering time, whereby only plants in categories 1, 2, and 3 experienced greater reproductive success if they started male flowering earlier (Table 1A; Figure 2C). Plants in the highest allocation category had similar reproductive success regardless of when they flowered (Table 1A; Figure 2C).

In contrast to our expectation that larger plants would sire more offspring, we discovered that neither plant height nor biomass had a significant effect on the number of seeds sired. Instead, we found that wider plants had greater male reproductive success (Table 1; *β* ± SE =0.08 ± 0.005; Figure S3). Plants located on the edge of the arrays sired significantly fewer seeds than plants located in the interior (edge: mean: 4.97 seeds, 95% CI: 4.06-6.08; interior: mean: 6.52 seeds, 95% CI: 5.45-7.82; Table 1).

### Mate diversity

There was a significant positive linear Bateman gradient indicating a strong relation between mating and reproductive success (Figure 3; *β* ± SE =0.71 ± 0.04, !*^2^=* 235.41, *P* <0.0001). We also found a significant quadratic term (*γ* ± SE =0.12 ± 0.03, !*^2^=* 24.75, *P* <0.0001). These relations were independent of male allocation.

**Figure 3.**
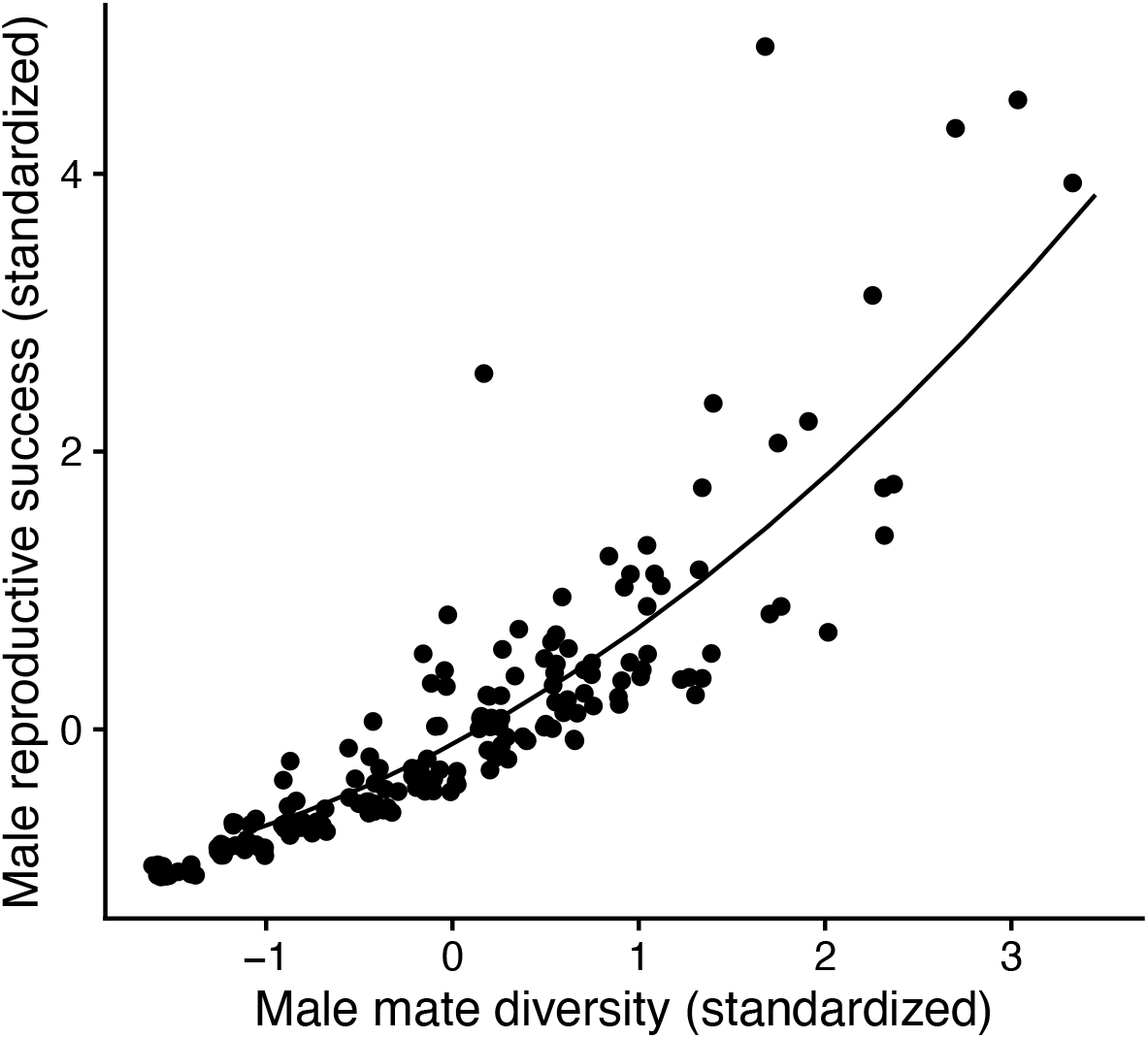
Male-specific Bateman gradient in *Ambrosia artemisiifolia* in experimental array conditions. The equation of best fit line is: *y*= 0.71*x* + 0.12*x*^2^-0.1, see text for statistical details.

We calculated male mate diversity by quantifying the number of mates to which a pollen-donor plant successfully dispersed pollen and sired seed. Of the genotyped seed, plants donated pollen to and mated with a mean=4.91 (SD=2.55, Range=1-14) different individuals. In raw numbers, males with larger male allocation had greater mate diversity (mean cat1: 3.71; cat2: 4.08; cat3: 5.61; cat4: 5.98), however this was likely a consequence of siring more seeds overall. In models that accounted for the total number of seeds sired, we found no significant effect of male allocation on mate diversity (Table 2).

**Table 2.** Models of effects on proportional mate diversity in Ambrosia artemiisifolia. Results for (A) male mate diversity model with factorial measure of male allocation, (B) male mate diversity model with continuous measure of male allocation (male inflorescence weight), and (C) female mate diversity. Models in A) and B) used a binomial distribution weighted by the number of seeds sired per individual, model C) used a poisson distribution and an offset parameter of the number of seeds genotyped per individual. Random effects (not shown) in all models included the effects of source population and block. Bold P-values indicate significant effects.

|  | **Term Condition** | **df** | **$\chi^2$** | ***P*-value** |
| --- | --- | --- | --- | --- |
| A) | Allocation category | 3 | 0.78 | 0.85 |
|  | Male flowering time | 1 | 9.37 | **<0.01** |
|  | Edge | 1 | 0.03 | 0.86 |
| B) | Male inflorescence weight | 1 | 0.01 | 0.91 |
|  | Male flowering time | 1 | 8.96 | **<0.01** |
|  | Edge | 1 | 0.09 | 0.76 |
| C) | Allocation category | 1 | 1.08 | 0.78 |
|  | Female flowering time | 1 | 8.93 | **<0.01** |
|  | Germination probability | 1 | 1.59 | 0.21 |

We previously reported that early flowering plants sired more total seeds (Figure 2C), when we investigate mate diversity it appears that many of those seeds were more likely to be full-sibs (i.e. sired on the same plant). Individuals that flowered later had a probability of greater mate diversity (Table 2; Figure 4A), that is, they donated pollen to and sired seeds on more unique mates. Amongst the morphological characteristics we measured (plant width, plant height, and biomass) none of them had a significant effect on mate diversity, after controlling for the number of seeds sired.

**Figure 4.**
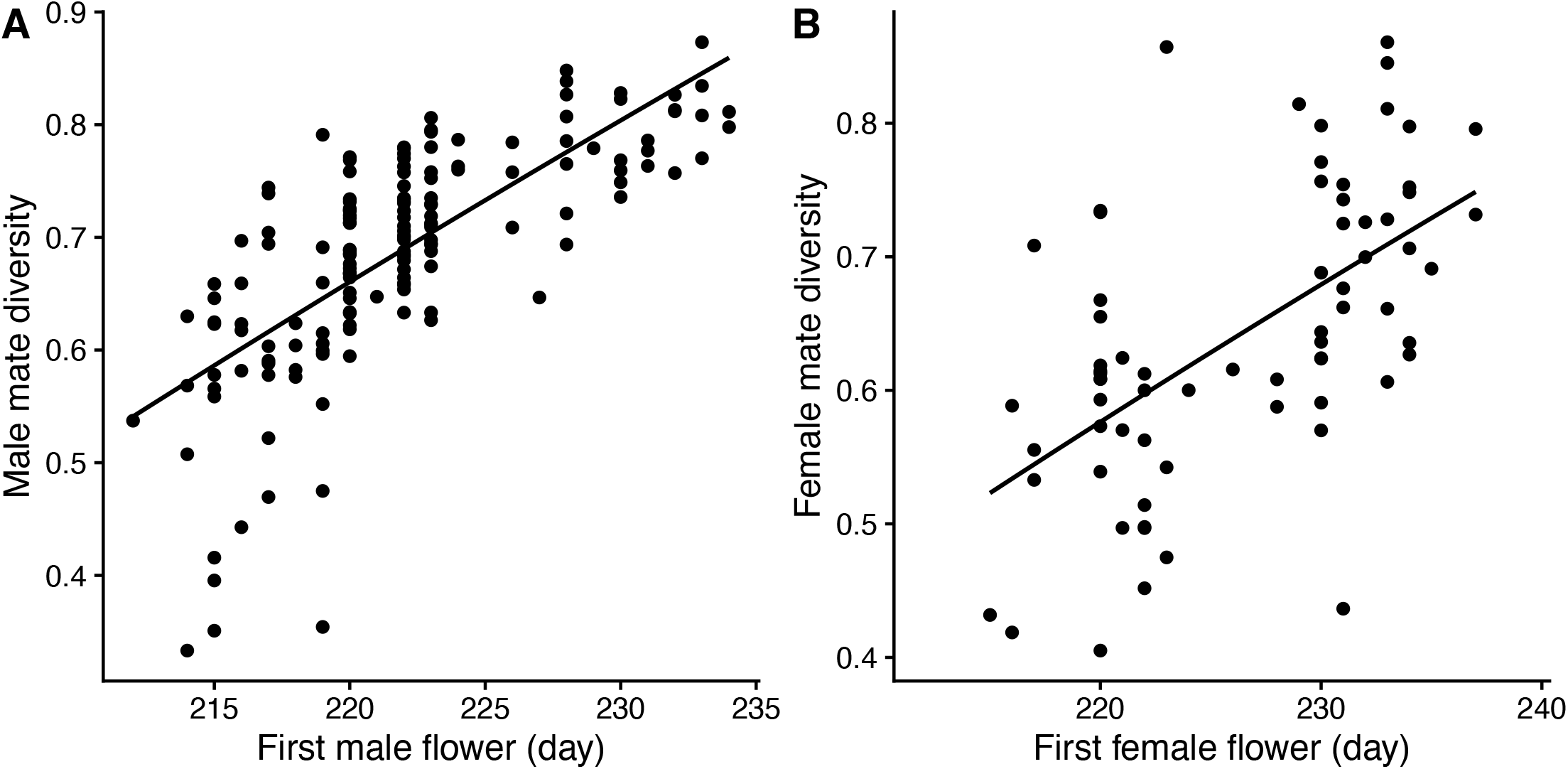
Effect of the Julian day of the start of (A) male flowering time and (B) female flowering time on male and female mate diversity (number of unique mating partners through female function), respectively, in *Ambrosia artemiisfolia* plants. Results represent model-adjusted and back-transformed values from ln estimates. See Table 2 for statistical details.

We quantified female mate diversity as the number of different pollen donor plants that sired seed on a given plant. We genotyped a subset of seed per plant (mean=17.25, SD=2.43, Range=11-22). Of these genotyped seed, plants mated with a mean=11.06 (SD=2.64, Range=5-16) different pollen donors. Plants that initiated female flowering later produced seeds that were sired by a greater diversity of pollen donors, than those with an earlier onset to female flowering (Table 2, Figure 4B). There was no association between female mate diversity and the allocation class of the individual, nor any of the measured morphological traits (height, width, biomass), nor germination probability.

## Discussion

Our results are consistent with the hypothesis that wind pollinated plants experience linear fitness gains with increased male allocation. Experimentally manipulating the number of male flowers elicited predicted changes in the number of offspring sired—plants with more male flowers sired significantly more offspring than plants with fewer male flowers. We found minimal direct effects of plant height on reproductive success, although plants with greater width (longer branches) had higher male fitness. We found significant, and surprising, influences of the timing of flowering, where an earlier onset to male flowering was beneficial for reproductive success, especially for plants with fewer male flowers. In contrast, individuals with later male flowering had relatively higher male mate diversity, and similarly later onset to female flowering was associated with seeds sired by a greater diversity of mates. Finally, our quantification of seed siring distance identified a strongly skewed and heavy-tailed distribution of successful pollen, with most seeds sired on nearest neighbours. Below, we consider the implications of these effects of sex allocation, intermate distance, and flowering time and their possible consequences on the evolution of reproductive strategies in wind pollinated plants.

### Incidental and unintended experimental consequences

Here we address the possible implications of several features of our experiment. First, we artificially created classes of plants with different male allocation to avoid confounding our intended variable of interest with overall condition or budget effects of plant size. This treatment necessarily involved removing male flowers, and for obvious reasons we could not artificially add male flowers. One implication of this is that we cannot test whether further exaggeration of male allocation will result in a saturation of reproductive success. We followed previous workers and fit a power function to our male gain curve data (Johnson and Yund 2009), and an implicit assumption of this is that males could theoretically produce enough pollen to monopolize fertilizations, which is presumably biologically impossible under at least some range of conditions. Second, because of the inflorescence architecture of *A. artemisiifolia*, when we removed male inflorescences we necessarily interfered with the height and width of plants, because male inflorescences are borne on terminal racemes. There was a modest, but significant, reduction in height and width for the lowest allocation treatment, but none of the other treatments differed significantly from each other (Figure S2). Our analyses accounted for this statistically by initially including all effects as covariates, but nonetheless we may have reduced power to identify direct effects of size on reproductive success. Third, we used a square lattice design which meant that plants are not equidistant to their first-order neighbours (i.e. horizontal and vertical neighbours are 0.5 m, but neighbours along the diagonal are 0.7 m). This is not a problem, per se, but should be considered when interpreting intermate distance.

### Male reproductive success and pollen dispersal

Our results are consistent with predictions from sex allocation theory that male fitness increases linearly with male allocation. To our knowledge, the only other explicit test of the male gain curve in wind pollinated plants comes from a natural stand of white spruce trees, where trees that produced more male cones sired a greater proportion of seeds (Schoen and Stewart 1986). There is some evidence that in the wind-pollinated herb *Mercurialis annua* plants with greater pollen production have greater reproductive success at high density (Tonnabel et al. 2019a). In this species, males with morphological traits that facilitated farther pollen dispersal achieved greater fitness, and only marginally improved fitness with greater pollen production. A previous study in *A. artemisiifolia* demonstrated that male reproductive success increased with height and male reproductive investment (Nakahara et al. 2018), but whether this was due to budget or direct effects of plant size was unclear. Other studies have reported that male reproductive success increases linearly with male allocation in spermcasting marine invertebrates (Yund and McCartney 1994; McCartney 1997), or shown that the male gain curve becomes more linear under certain ecological conditions like intense local sperm competition or mating over larger distances (Yund 1998; Johnson and Yund 2009). Our finding of a linear gain curve means that increased allocation to pollen production should be favored under some ecological conditions. Previous work demonstrated substantial genetic variation for plasticity in male allocation in ragweed (Friedman and Barrett 2011a), so that allocation may respond adaptively to ecological conditions.

Because we experimentally manipulated plant size, we removed budget effects (i.e. the effect where plants in better condition or larger plants invest more in reproduction because they have a larger resource budget), any remaining effects of plant size on male fitness would more likely be due to direct effects of size (Klinkhamer et al. 1997). Experimentally controlling for male investment, we found no association between plant height and siring success. Similarly, there was no significant effect of plant height on male reproductive success in *M. annua* (Tonnabel et al. 2019). But our result is discordant with a different study in *A. artemisiifolia* that found a weak effect of plant height on siring success, although the effect depended on model assumptions about neighbourhood size (Nakahara *et al*. 2018). All of these studies were conducted in common garden arrays where the benefit of height might be diminished compared to natural settings, because of the reduction in intervening vegetation. Nonetheless. for herbaceous plants occurring in open environments, siring success may not be strongly influenced by plant height, but by other aspects of size that affects the ability to better disperse pollen.

We identified a benefit of plant width on male reproductive success. Although we did not initially predict this, the architecture of the plant suggests that this effect is due to the aerodynamics of pollen release in wind pollinated plants (Niklas 1985). All objects are surrounded by a layer of still air, and the size of the boundary layer is determined by the size of the solid structure disrupting air flow. An important adaptation for wind pollinated plants is positioning male flowers away from vegetative structures to get them out of the boundary layer and enhance pollen liberation from anthers (Timerman and Barrett 2021). In many species this involves extending the anthers on long filaments that vibrate in the wind (Timerman and Barrett 2018). However, in *A. artemisiifolia* dehiscent anthers extend only just below the downward-pointing floret (Payne 1963), and so the position of the staminate head on branches that extend beyond foliage may better expose them to the wind to increase vibration and facilitate the release of pollen into the airstream (Friedman and Harder 2005). Similarly, in *M. annua* selection favors males with wider diameters in combination with longer branches and greater biomass (Tonnabel et al. 2019a). Thus, there is increasing evidence that branching architecture may provide direct beneficial effects for male fitness in wind-pollinated plants.

Wind-dispersed pollen typically has a leptokurtic distribution from point sources (Bateman 1947; Gleaves 1973; Levin and Kerster 1974), so that the seed set of recipients should decrease rapidly with distance from the pollen donor. Our results are consistent with this. Most plants sire seeds on their nearest neighbours, and the distance between mates shows a strongly fat-tailed distribution. Other studies have found similar results, for example pollination success in *Taxus* declined with plant spacing (Allison 1990); in dioecious *Thalictrum* species female plants at greater distance from males had reduced seed set (Steven and Waller 2007); and in *Festuca pratensis*, most pollen was deposited within 75m of donors (Rognli *et al*. 2000). In our experiment, none of the morphological variables we measured had any significant effect on the distance between mates. Similarly, Nakahara et al. (2018) found no effect of plant height on the the maximum distance between mates. These findings are in contradiction to the theoretical expectation that taller wind pollinated plants will have farther pollen dispersal (Burd and Allen 1988), and indicate that plant height may have only limited consequences for pollen dispersal in herbaceous plants in open fields. Indeed, successful matings at the farthest distances in our experiment (between blocks) appear to be stochastic. Nonetheless, for plants that sired fewer seeds overall, these long-distance siring events represent a greater proportion of their total seeds sired. While most mating occurs very locally in wind pollinated plants, rare longer-distance mating events may profoundly impact genetic structure and patterns of genetic variation (Loveless and Hamrick 1984).

### Mating portfolios and the benefits of mate diversity

We found high variance in both male reproductive success and male mating success, as expected under Bateman’s principle and sexual selection (Tonnabel et al. 2019b). To investigate the fitness accrued by mating with more partners, we estimated the Bateman gradient for male function, and identified significant positive linear and quadratic (accelerating) terms. This relation was not influenced by allocation category or male inflorescence weight, suggesting consistent benefits for all individuals. Similarly, while individuals with more male flowers had greater mate diversity in absolute terms, this was entirely driven by their concomitant increase in reproductive success. This finding supports the proposition that multiple mating in plants is a by- product of selection on male function to increase siring success (Pannell and Labouche 2013). Mating with multiple female partners (high mate diversity) provides a quantitative advantage through male function, and our data corroborate that mating opportunities constrain male reproductive success.

There is scant evidence that mate diversity, per se, is under selection or beneficial (Barrett and Harder 2017). However, several lines of evidence suggest that genetic diversity may be advantageous through a process of ‘genetic bet-hedging’. When stochasticity is incorporated into measures of natural selection, then fitness depends on both the mean and the variance in offspring number (Gillespie 1974) and increasing the variance in offspring number of a genotype will decrease its fitness (Gillespie 1977). To the extent that mating between any two individuals leads to low fitness (e.g. due to genetic compatibility), then having a greater diversity of mates could reduce the variance in offspring number and increase mean fitness. A second related argument is that genetic diversity within families is beneficial for offspring success in the face of temporal or spatial heterogeneity. When environments are heterogeneous, offspring diversity raises the chances that some offspring succeed, thus decreasing the variance in success and increasing geometric mean fitness (Slatkin 1974; Simons 2011). In animals, various lines of evidence suggest that genetic bet hedging is unlikely to be solely responsible for maintaining polyandry, unless the costs of multiple mating are very low (Yasui 1998; Jennions and Petrie 2000). Indeed, in plants the costs of multiple mating are likely low, especially for wind pollinated plants that are not investing in showy flowers, raising the likelihood that genetic bet- hedging provides a selective advantage.

Mate diversity can also be considered from the perspective of the interactions among siblings – both during seed development and subsequent dispersal and establishment. First, genetic variation among developing embryos provides an opportunity for maternal resources to be distributed to the highest quality embryos and potential abortion of incompatible or low-quality embryos (Zeh and Zeh 1996; Haig and Westoby 1988). However, sibling competition within the developing fruit can be detrimental to both maternal and paternal parents, and several plant reproductive strategies may have evolved to reduce mate diversity within an ovary (Kress 1981; Bawa 2016). For example, generalist, indiscriminate pollinators are most likely to deliver unrelated pollen grains onto stigmas, and there is a well-established association between uni- ovulate flowers and abiotic and generalist pollinators (Charlesworth 1993; Friedman and Barrett 2011b). Like most wind pollinated plants. *A. artemisiifolia* has flowers with single ovules, so differential maternal investment would occur among developing fruits on a plant. Second, greater mate diversity increases the number of half-sib families rather than full-sib families, so in species with restricted seed dispersal establishing seedlings are less related. Genetic diversity among seedlings reduces sib-competition thereby increasing maternal (and paternal) fitness (Cheplick 1992). Furthermore, to the extent that pollen dispersal is limited, genetic diversity of offspring will reduce biparental inbreeding (Uyenoyama 1986) and any accompanying inbreeding depression (Charlesworth and Willis 2009). Future work is necessary to partition the relative influence of these factors on selection for mate diversity.

### The benefits of temporal separation in flowering

The temporal separation of flowering between sexes (dichogamy) generates uneven sex ratios across a flowering season (Brunet and Charlesworth 1995; Sargent and Roitberg 2000), which is paradoxical because frequency-dependent selection should act to equalize the availability of pollen and ovules at every point in time. The benefits of dichogamy (and protandry in particular) include avoiding interference between male and female sex organs (Lloyd and Webb 1986; Bertin 1993) and preventing selfing (Darwin 1876). Neither of these mechanisms are likely responsible for protandry in *A. artemisiifolia* because the species is monoecious and self-incompatible. In a simulation model of the asynchrony in timing of pollen and ovule presentation, Medan and Bartoloni (1998) demonstrated that selection favours protandrous genotypes when there is substantial overlap in male and female function; as they have the greatest access to mates (both pollen and ovules), although flowering too early can be wasteful.

Several results support this balance of influences on protandry. We found that earlier onset to male function increased male reproductive success (especially for plants that had lower overall male allocation), but later flowering increased the probability of mating with diverse partners. Although we cannot identify the mechanisms here, we speculate that the potential for male mating success depends on competition for access to ovules from other individuals—more ovules will be available later in the season, but there will also be greater pollen competition. Plants benefit from early flowering by monopolizing siring opportunities on the few individuals that have female flowers, but shifting male flowering to later when more plants are blooming benefits relative mate diversity. Similarly, slightly delaying female function resulted in seed sired by a more diverse pollen pool. Other mechanisms may be at play, for example if plants with earlier flowering donated their pollen to more fecund mates, had greater pollen competitive ability, or if their mates had declining fruit set (Brunet 1996, Weis and Kossler 2004, Austen and Weis 2016b). The pattern in our results suggests that the staggered onset to male and female flowering benefits both sex functions (in “harmony” between the sexes: Delph and Ashman 2006), while fecundity selection and sexual selection through male function are acting in opposing directions on the start of male flowering.

Sex ratio selection and availability of ovules will eventually constrain the continued evolution of dichogamy, demonstrated in a theoretical model by Sargent et al. (2006). An extended blooming season alleviates some of the costs of skewed sex ratios—the earliest blooming plants sacrifice some male mating opportunities that are lost due to an absence of available ovules, but the wasted pollen represents a small fraction of overall investment. This scenario is exemplified by *A. artemisiifolia*, where flower production increases, and under some conditions accelerates, through time (Friedman and Barrett 2011a: see Figure 4), mitigating the costs of wasted mating opportunities by early blooming flowers. Further, plants flower for 6 weeks or more, and the average degree of dichogamy in our experiment was 4.2 days (range= -9 to 17 days; 9% of plants were protogynous), so plants express both sex functions for the majority of their flowering.

The duration of overlap between male and female function, or their temporal separation, may be an adaptive response to environmental stochasticity. Under conditions where resource availability is unpredictable and resource acquisition varies during the flowering season, the best allocation strategy should be in favor of the sex function with the higher return on investment (Zhang 2006). Indeed, *Ambrosia artemisiifolia* experiences substantial plasticity in sex allocation and in the degree and order of dichogamy (Paquin and Aarssen 2004; Friedman and Barrett 2011a). Here we have demonstrated that the male gain curve is linear or nearly so, and if we assume that the female gain curve is mostly linear (Nakahara et al. 2018), then we would expect a sharp transition between sex functions (Zhang et al. 2006). The gradual transition that we observe may be explained because male flowers are photosynthetic and contribute to available resources (Bazzaz and Carlson 1979), and when reproductive resources are not a pool, but an income, then the constraints on staminate flowering are altered (Burd and Head 1992).

Nonetheless, although the male gain curve is linear, there are likely diminishing marginal returns of male investment through time due to the saturation of ovules.

## Conclusion

Experimental tests of the shape of male gain curves and factors that affect it are scarce compared to the amount of existing theoretical work. Our study is one of the few studies that empirically test theoretical expectations of male gain curve in wind pollinated plants. In agreement with predictions, we found a linear increase in fitness returns for increasing investment in male function. Fitness through male function is likely limited by the availability of mates, and increasing male investment results in proportional increases in the number of mating partners. In wind-pollinated *A. artemisiifolia* there is likely strong selection on producing more pollen, with the direct benefit increasing siring success and indirectly leading to more mating partners. While early onset male flowering (particularly for lower male allocation plants) benefits male reproductive success, later onset results in greater probability of mating with diverse partners through both male and female function. Together this suggest an adaptive role for protandry to adjust the pool of competing pollen for available ovules, and a balance between fecundity selection and sexual selection through male function. Plants with lower male allocation might experience particular benefits by flowering earlier and avoiding the competitive arena during full population blooming, although a more explicit test of this prediction is necessary to rule out alternative explanations.

## Author Contributions

JF conceived the study, AA and JF designed the study, AA conducted the field work and genetic assays, AA and JF analyzed the data and JF drafted the manuscript with input from AA.

## Acknowledgements

The authors acknowledge funding from NSF EAGER 1546106 to JF, a R.C. Lewontin Early Award from the Society for the Study of Evolution to AA, and support from Syracuse University and Queen’s University. Thanks to Matthew Rubin and Karine Leydet for assistance with field and laboratory work. The ideas and analyses in this manuscript were greatly improved by helpful discussions with Spencer Barrett, David Timerman, Scott Pitnick, and Crispin Jordan.

## Supplementary Materials

**Figure S1.**
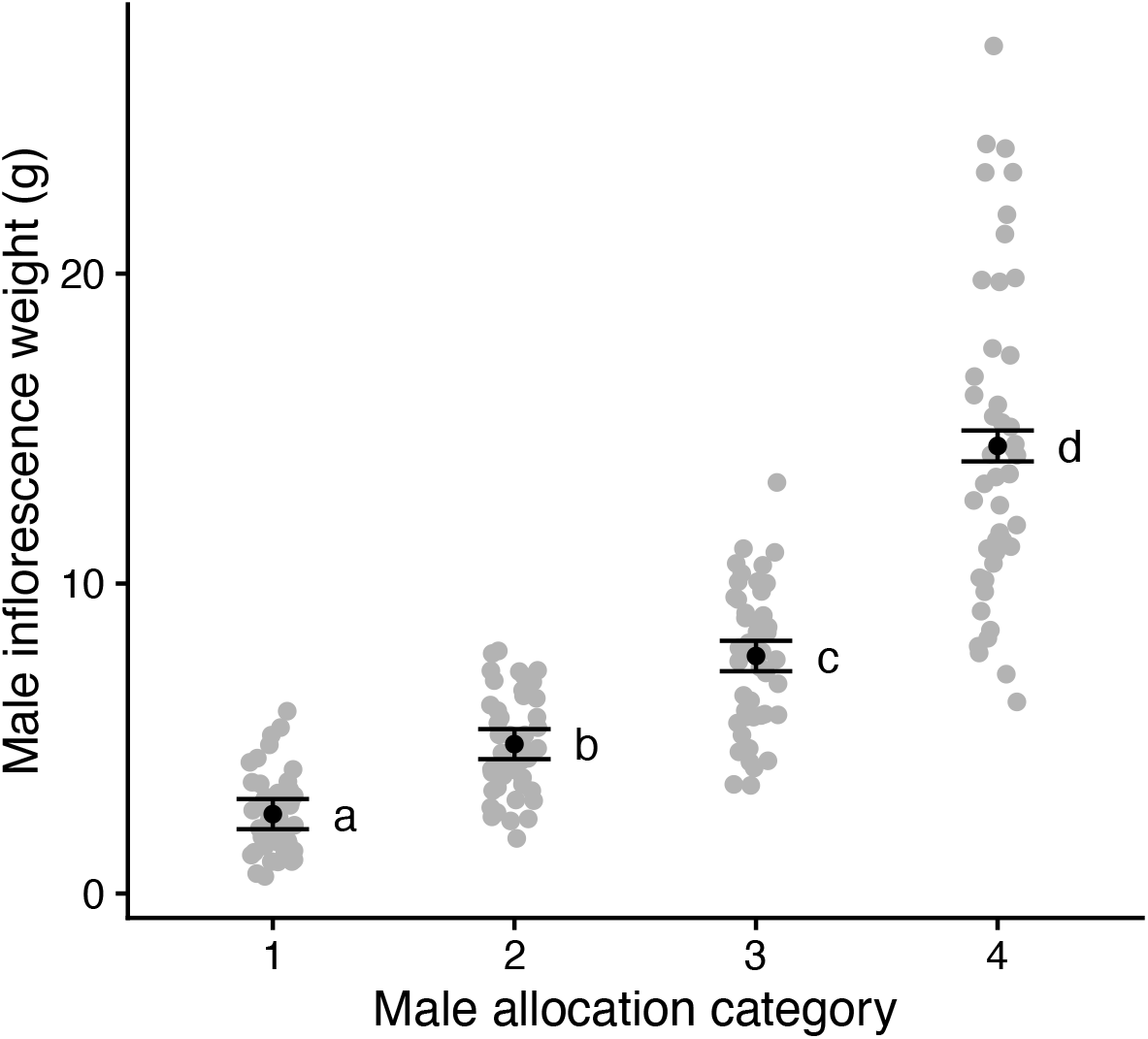
Effects of the categorical experimental treatment of male flower manipulation on mean (±S.E.) male inflorescence weight in experimental *Ambrosia artemisiifolia* plants. Raw data are plotted, and means associated with different lowercase letters differed significantly.

**Figure S2.**
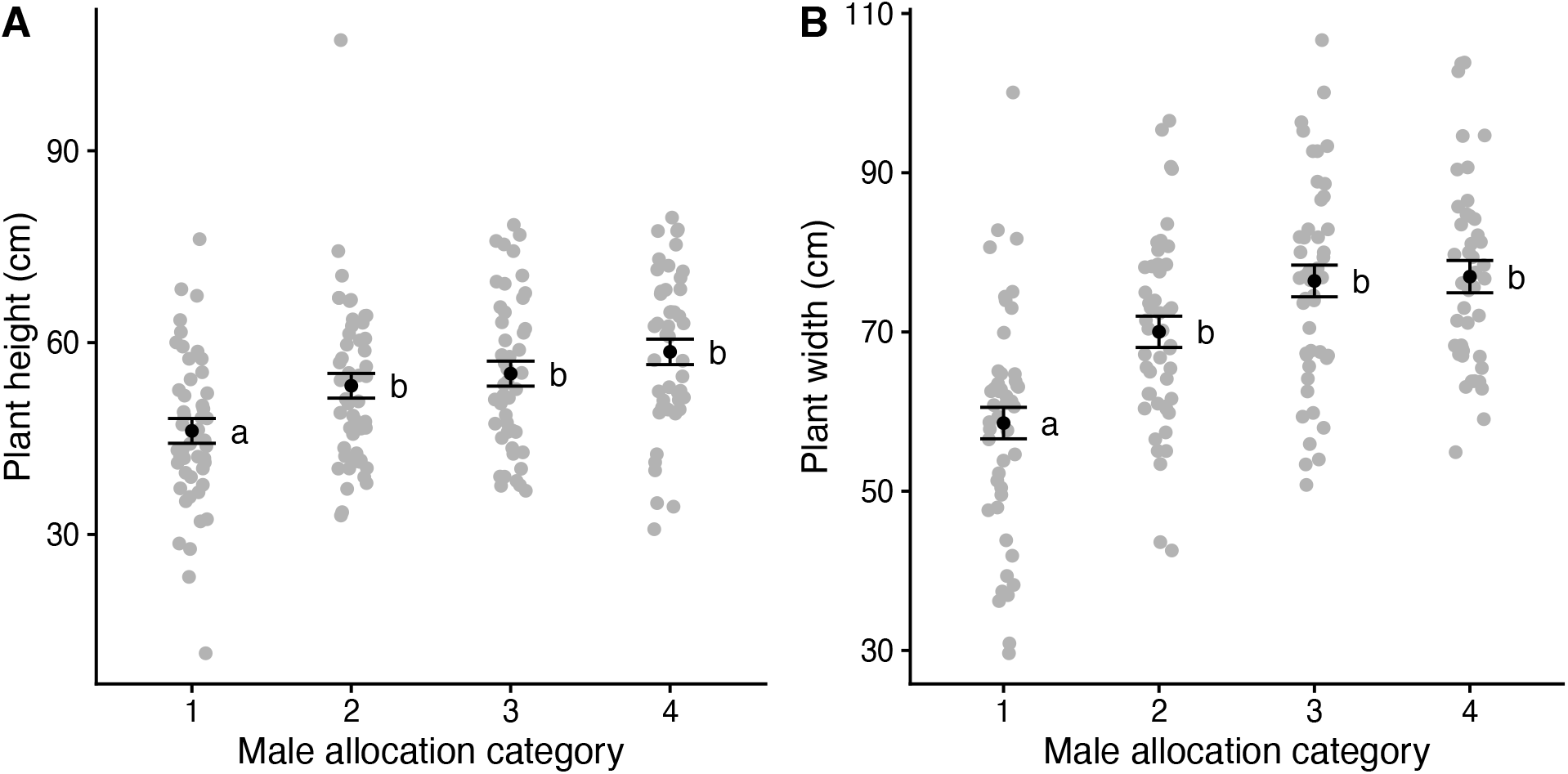
Effects of the categorical experimental treatment of male flower manipulation on mean (±S.E.) A) plant height, and B) plant width, in experimental *Ambrosia artemisiifolia* plants. Raw data are plotted, and means associated with different lowercase letters differed significantly.

**Figure S3.**
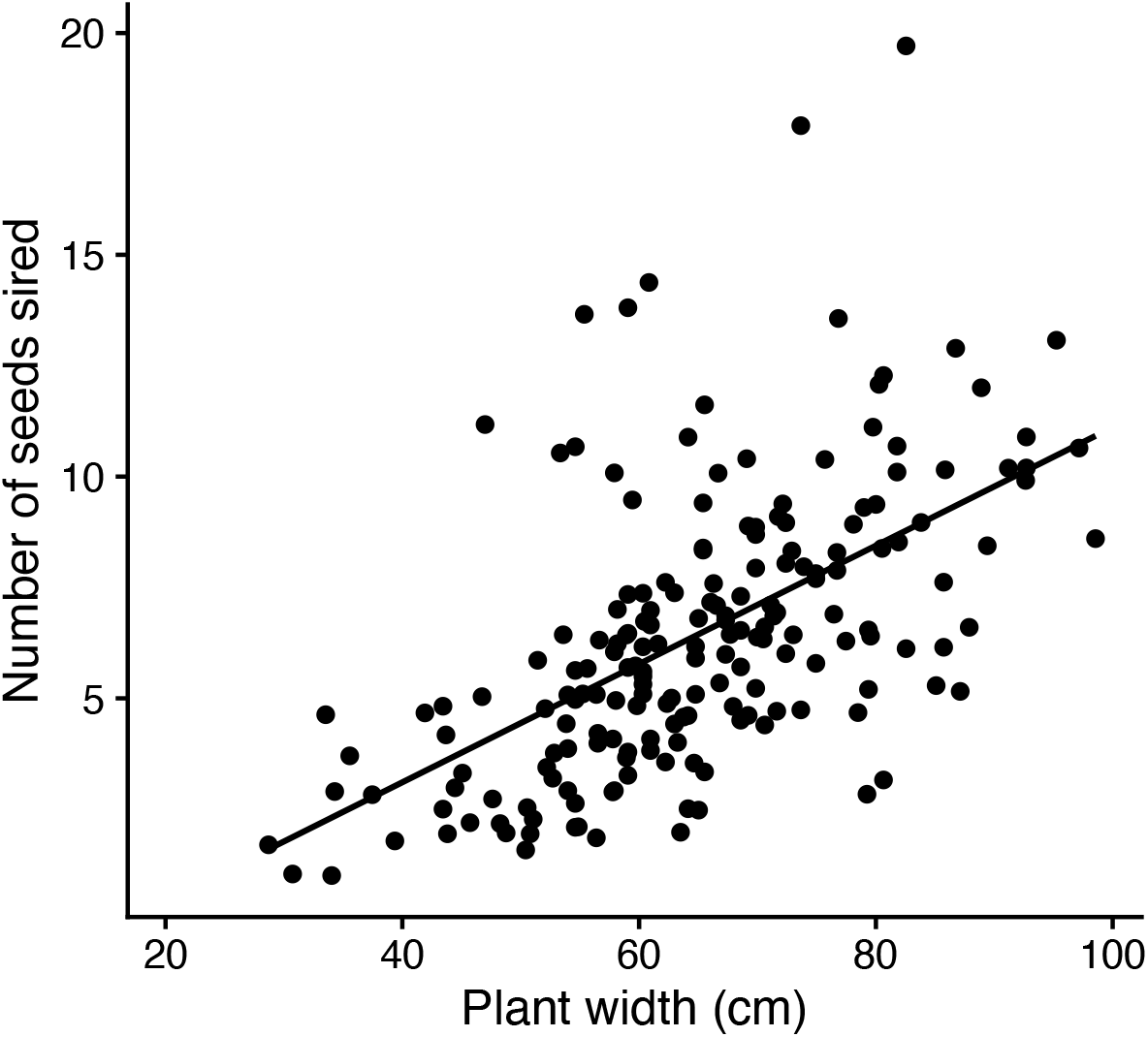
Influence of plant width (cm) on the number of seeds sired (male reproductive success) in *Ambrosia artemiisfolia* plants. Results represent model-adjusted and back-transformed values from ln estimates. See Table 1 for statistical details.

**Table S1.**
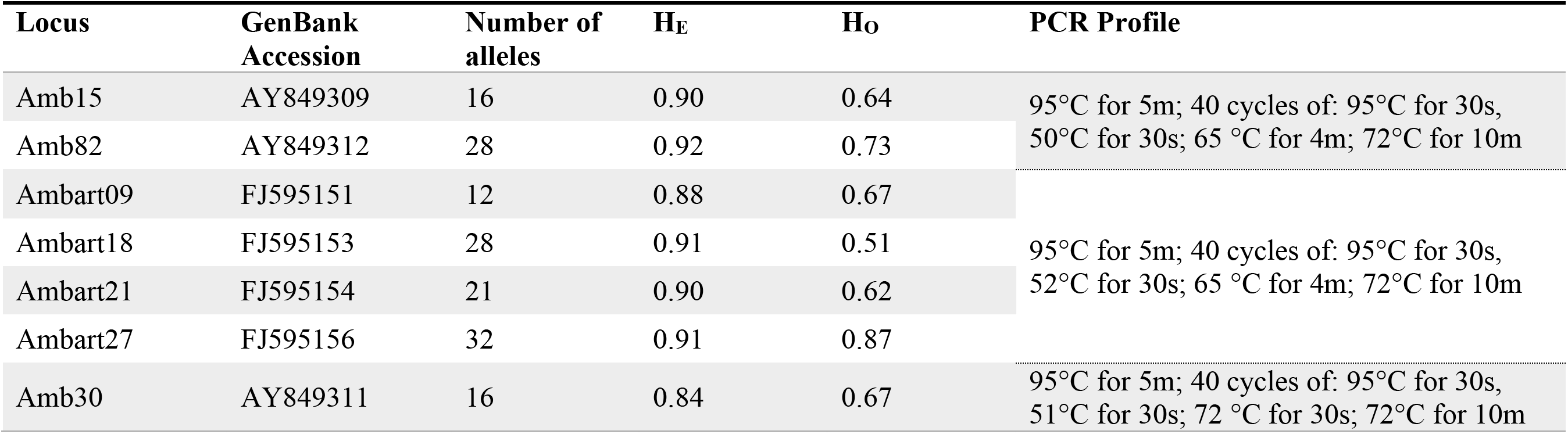
Characteristics of the seven microsatellite loci used for paternity analysis. GenBank Accession, number of alleles, expected heterozygosity (H_E_), observed heterozygosity (H_O_), and PCR conditions for amplification are provided. All PCR reactions included a final volume of 10µl containing approximately 10-20ng of genomic DNA, 2µl of 5X Taq buffer, 0.5µl of 25mM MgCl_2_, 1µl of 0.25mM of each dNTP, 0.1µl of 10µM of each forward and reverse primers and 0.5U of Taq Polymerase.

